# Comparison of *Galleria mellonella*, Epithelial Cell Cytotoxicity, and Mouse Model of Bacteremia to Measure *Pseudomonas aeruginosa* Virulence

**DOI:** 10.64898/2026.03.16.712016

**Authors:** Aliki Valdes, Christopher Axline, Travis J. Kochan, Sophia Nozick, Timothy Ward, Issay Niki, Ethan VanGosen, David Hynes, Julia Nelson, Preeti Garai, Tania Afzal, Daniel Amusin, Sumitra D. Mitra, Timothy L. Turner, William Cheng, Joanne J. Lee, Prarthana Prashanth, Nathan B. Pincus, Jonathan P. Allen, Jake Hauser, Egon A. Ozer, Kelly E. R. Bachta, Cheng-Hsun Chiu, Antonio Oliver, Alan R. Hauser

## Abstract

Considerable effort has focused on identifying alternatives to mouse models in research studies. In the field of bacterial pathogenesis, *Galleria mellonella* and epithelial cell lines have been widely used for this purpose, but the concordance of these models with mice remains unclear. To begin to address this knowledge gap, we used 105 clinical isolates of *Pseudomonas aeruginosa* for which the virulence had been previously determined in a mouse bacteremia model. A semistrong correlation was observed between *G. mellonella* median time to 50% mortality and mouse 50% pre-lethal dose (LD_50_) values (Spearman’s rank correlation coefficient [ρ] = 0.75), whereas percent A549 epithelial-like cell lysis during co-culture showed a weak correlation to mouse LD_50_ values (ρ = -0.47). Given the stronger correlation between *G. mellonella* and mouse virulence, we next examined whether *G. mellonella* could substitute for mice when asking questions about the virulence of large numbers of *P. aeruginosa* isolates. Results from mice indicated that isolates with resistance to more antibiotics were significantly less virulent, and the use of *G. mellonella* identified the same inverse correlation. Furthermore, both models found no evidence for the existence of hypervirulent clonal lineages. In particular, isolates belonging to sequence types defined as high-risk clones were not consistently more virulent than other isolates, despite the known association of high-risk clones with poor clinical outcomes. These findings suggest that *G. mellonella* can serve as an adequate substitute for mice when addressing specific population-based virulence questions, although conclusions should be confirmed in mice.

**Author Summary:** We found that virulence measurements in a *G. mellonella* infection model showed a semistrong correlation with those from a mouse bacteremia model and that this insect larval model adequately detected population-level trends similarly to mice. In contrast, A549 epithelial-like cell lysis during bacterial co-culture correlated less well with mouse virulence. Together, these results support the use of *G. mellonella* as a scalable, low-cost, and humane first-line model for assessing *P. aeruginosa* virulence but also indicate that conclusions should be validated in mice.

## Introduction

Mouse infection models are arguably the gold standard for studying the virulence of bacteria such as *Pseudomonas aeruginosa* [1] and have many advantages, such as human-like innate and adaptive immune systems, genetically modifiable genomes, and an extensive body of comparative literature. However, mouse infection models are costly, highly regulated, and ethically controversial, which limits their utility in high-throughput assays. *Galleria mellonella* and mammalian cell culture models circumvent many of these disadvantages [2,3]. Both may be maintained at 37°C, the mammalian body temperature at which some *P. aeruginosa* virulence genes are upregulated [4]. Cell co-culture assays have the added advantage of allowing for use of human cells lines, whereas *G. mellonella* harbor hemocytes that function similarly to mammalian neutrophils [2,5], the immune cell primarily responsible for containing *P. aeruginosa* infections in mice and people [6–9]. As a result, both *G. mellonella* and cell culture models have been widely used to quantify *P. aeruginosa* strain virulence [10–17]. However, the degree to which these models mirror *P. aeruginosa* infection outcomes in mice has been examined in only a few small studies [10–14].

Most *P. aeruginosa* isolates fall into one of two genetically distinct clades: Group A and Group B [18,19]. These clades are largely distinguished by the presence of either the *exoS* or *exoU* gene, respectively, which encode type III secretion system (T3SS) effectors. The presence of the *exoU* gene and secretion of its corresponding effector protein are associated with more severe disease [15,20–23]. Within each of these large clades, subclades of multilocus sequence types (STs) exist, which are defined as isolates with the same alleles across seven housekeeping genes [24]. Some STs have been designated as high-risk clones (HRCs) because of their global distribution, the disproportionate number of infections they cause, their high degree of antibiotic resistance, and their association with poor clinical outcomes (27, 28). Even within a single *P. aeruginosa* ST, substantial isolate-to-isolate genomic variability exists, which drives phenotypic differences in virulence and antibiotic resistance [20,21,30–33]. Some antibiotic resistance mutations confer fitness costs that reduce virulence [27,34–37], whereas others do not [37–39]. Clarifying strain-to-strain differences in virulence and antibiotic resistance and the interplay between the two is crucial to a deeper understanding of the pathogenesis of *P. aeruginosa* infections.

In this study, we used 105 *P. aeruginosa* clinical isolates to examine how virulence measurements from the *G. mellonella* and mammalian cell co-culture models compare to those from mice. Results from *G. mellonella* showed a semistrong correlation with those from the mouse model, whereas cell cytotoxicity results correlated less well. We then used the *G. mellonella* and mouse results to examine the relationships between *P. aeruginosa* virulence and antibiotic resistance and whether hypervirulent clades of *P. aeruginosa* exist.

## Materials and Methods

### *P. aeruginosa* isolates and culture conditions

A description of the *P. aeruginosa* isolates used in this study has been previously published [32,40]. Seventy-one isolates (“PABL”) were obtained at Northwestern Memorial Hospital in Chicago, IL, USA from 1999 to 2003 from adults with *P. aeruginosa* bacteremia [41]. Fifteen isolates (“PAC/S”) were obtained from the blood or peritoneal fluid of pediatric patients with Shanghai fever at Chang Gung Children’s Hospital in Taiwan from 2003 to 2008 [42]. Nineteen isolates (“PASP”) were collected from patients with bacteremia in Spain between 2008 and 2009 [43]. Bacteria were routinely cultured in lysogeny broth (LB) at 37°C with shaking at 250 rpm or on LB agar plates.

### Whole-genome sequencing and phylogenetic analysis

The whole-genome sequence of each isolate has been previously published [32,40,44]. Multilocus sequence typing (MLST) was performed using the *P. aeruginosa* PubMLST website (https://pubmlst.org/paeruginosa/)[45].

### Mouse infection model

A mouse model of bacteremia using tail vein injection of bacteria had been previously used to estimate the virulence of each *P. aeruginosa* isolate [32,40]. Briefly, each isolate was tested at a minimum of two doses, with 3-5 mice per dose (≥9 mice per isolate). The bacterial CFU required to cause pre-lethal illness in 50% of mice (LD_50_) following tail vein injection was calculated. Mice meeting pre-defined criteria [23] for pre-lethal illness were euthanized and scored as dead. For most of the isolates, LD_50_ values were obtained and reformatted from Table S3 of Pincus et al. [32], distributed under the terms of the Creative Commons Attribution 4.0 International license (https://creativecommons.org/licenses/by/4.0/). For PABL019, the mouse LD_50_ was obtained from Allen et al. [40].

### *G. mellonella* infection model

*G. mellonella* were infected as previously described [10]. A defined bacterial dose was injected into *G. mellonella* larvae, and infected larvae were monitored for mortality over time to calculate a 50% lethal time (LT_50_), at which half the infected larvae had died (Supplemental Fig. 1A). This process was then repeated with a range of doses. The natural logarithms of these doses were plotted against the corresponding LT_50_ values, and a linear regression was performed to establish the ln(CFU) vs. LT_50_ relationship (Supplemental Fig. 1B). This process was repeated for all isolates (Supplemental Data). To compare isolate virulence, regression equations were used to predict the LT_50_ values of each isolate at a dose of 3,000 CFU, the approximate median dose across all *G. mellonella* infections. This approach circumvents the need to experimentally prepare inocula of 3,000 CFU with high accuracy, which can be challenging for such a small dose and with a large collection of diverse isolates. Comparisons were also made at calculated doses of 300 and 30,000 CFU.

### Cytotoxicity assay

A549 cells (ATCC) were grown at 37°C in Dulbecco’s Modified Eagle Medium (Corning) supplemented with 10% fetal bovine serum (Corning) under 5% CO_2_ conditions. Cytotoxicity was quantified by measuring lactate dehydrogenase (LDH) release at 3 hours post-infection using the CyQUANT™ LDH Cytotoxicity Assay (Invitrogen) per the manufacturer instructions.

### Immunoblot analysis

For detection of *in vitro* secreted ExoU and ExoS proteins, a detailed procedure is provided in Supplemental Information. Immunoblot results for the PABL set of isolates were previously published [40].

### Antibiotic susceptibility

Minimum inhibitory concentrations (MICs) were performed using microbroth dilution assays as previously described [46]. Each isolate was tested against six agents: tobramycin (aminoglycoside), piperacillin/tazobactam (penicillin/β-lactamase inhibitor), aztreonam (monobactam), cefepime (cephalosporin), ciprofloxacin (fluoroquinolone), and meropenem (carbapenem). Antibiotic resistance was determined as non-susceptible (i.e., MICs in the intermediate or resistant range) or susceptible according to the Clinical and Laboratory Standards Institute 2024 breakpoints. Isolates were ranked from 0 (susceptible to all 6 antibiotics) to 6 (non-susceptible to all 6 antibiotics). All MIC tests were performed in duplicate; if the duplicates disagreed, a third replicate was performed.

### Statistical analyses

All statistics were performed using R (v4.5.1) or GraphPad Prism (v10.2.3). The statistical tests used are indicated in the corresponding figure legends. *P* ≤ 0.05 was considered significant. Spearman’s rank correlation coefficient (ρ) strength was interpreted using previously published references: weak (|ρ| < 0.5), moderate (0.5 ≤ |ρ| < 0.75), semistrong (0.75 ≤ |ρ| < 0.85), and strong (|ρ| ≥ 0.85)[47].

## Results

### Characterization of a collection of 105 *P. aeruginosa* clinical isolates

To compare the three models of infection, we utilized a collection of previously sequenced *P. aeruginosa* clinical isolates cultured from patients in the United States, Spain, and Taiwan (Fig. 1). These isolates were genetically diverse, consisting of 59 different STs (Supplemental Data). Both the Group A and Group B clades of the *P. aeruginosa* population structure were represented, with 64 isolates containing only the *exoS* gene (*exoS*^+^) and 38 containing only the *exoU* gene (*exoU*^+^). PABL031 contained both effector genes (*exoS*^+^/exo*U*^+^) while PABL071 and PABL043 contained neither (*exoS*^-^/exo*U*^-^). Immunoblot analyses indicated all but 20 isolates secreted detectable amounts of the corresponding T3SS effector protein under inducing *in vitro* growth conditions. Of these isolates (ExoS**^-^/**ExoU**^-^**), 14 were *exoS*^+^, 4 were *exoU*^+^, and 2 were *exoS*^-^/exo*U*^-^ (Supplemental Data). These findings show that the collection of *P. aeruginosa* isolates is phylogenetically diverse and represents a variety of type III effector genotypes and secretion phenotypes.

**Figure 1.**
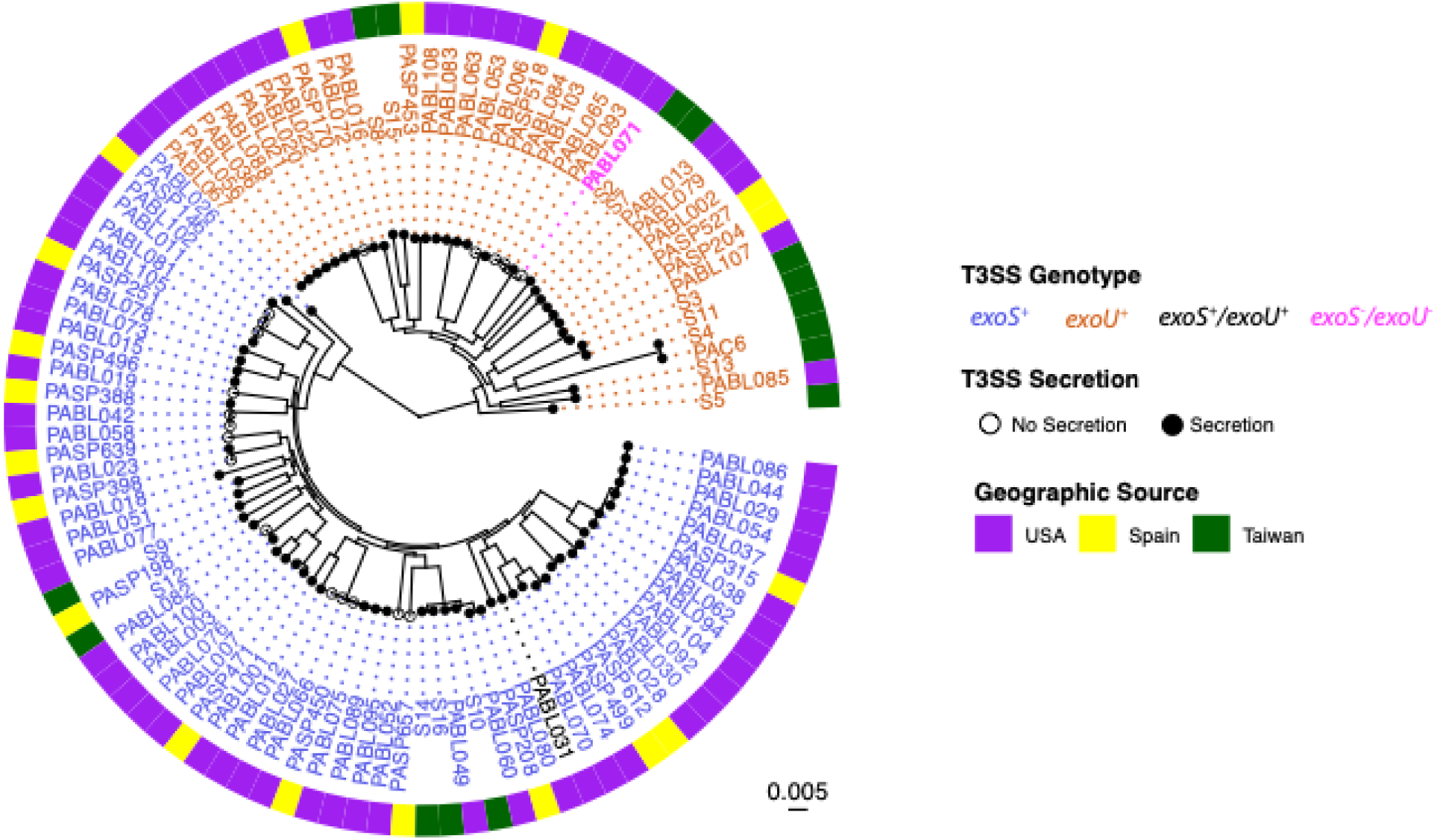
Phylogenetic and phenotypic diversity of *P. aeruginosa* clinical isolates. A maximum-likelihood core genome phylogenetic tree is shown, with a concentric ring indicating isolate geographic source. The presence of the *exoS* or *exoU* genes is indicated by the font color of the isolate name, and *in vitro* secretion of T3SS effectors ExoS and ExoU is indicated by tips with filled circles (secretion-positive) or open circles (secretion-negative). PABL043 (a PA7-like strain) is excluded from this tree because it is genetically highly divergent from the other isolates.

### Virulence in a mouse model of bacteremia

All isolates had been previously assessed for virulence in mice by measuring the 50% pre-lethal dose (LD_50_) in a tail vein injection bacteremia model [32,40]. These results are reproduced in Figure 2. The isolates varied markedly in their virulence, with LD_50_ values ranging over more than three logs (1000-fold) with a median LD_50_ value of 7.4 x 10^6^ (10^6.9^) CFU. PABL012, an ExoS^+^ isolate, was the most virulent, with an LD_50_ value of 6.3 x 10^5^ (10^5.8^) CFU (Supplemental Data). PABL019, an ExoS^-^ isolate that was obtained from the bloodstream of a patient with cystic fibrosis (CF), had the highest LD_50_ value of 1.3 x 10^9^ (10^9.1^) CFU. These findings demonstrate substantial differences in the capacity of individual *P. aeruginosa* isolates to cause severe infections. They also serve as a reference for measuring the performance of the *G. mellonella* and cell co-culture models of infection.

**Figure 2.**
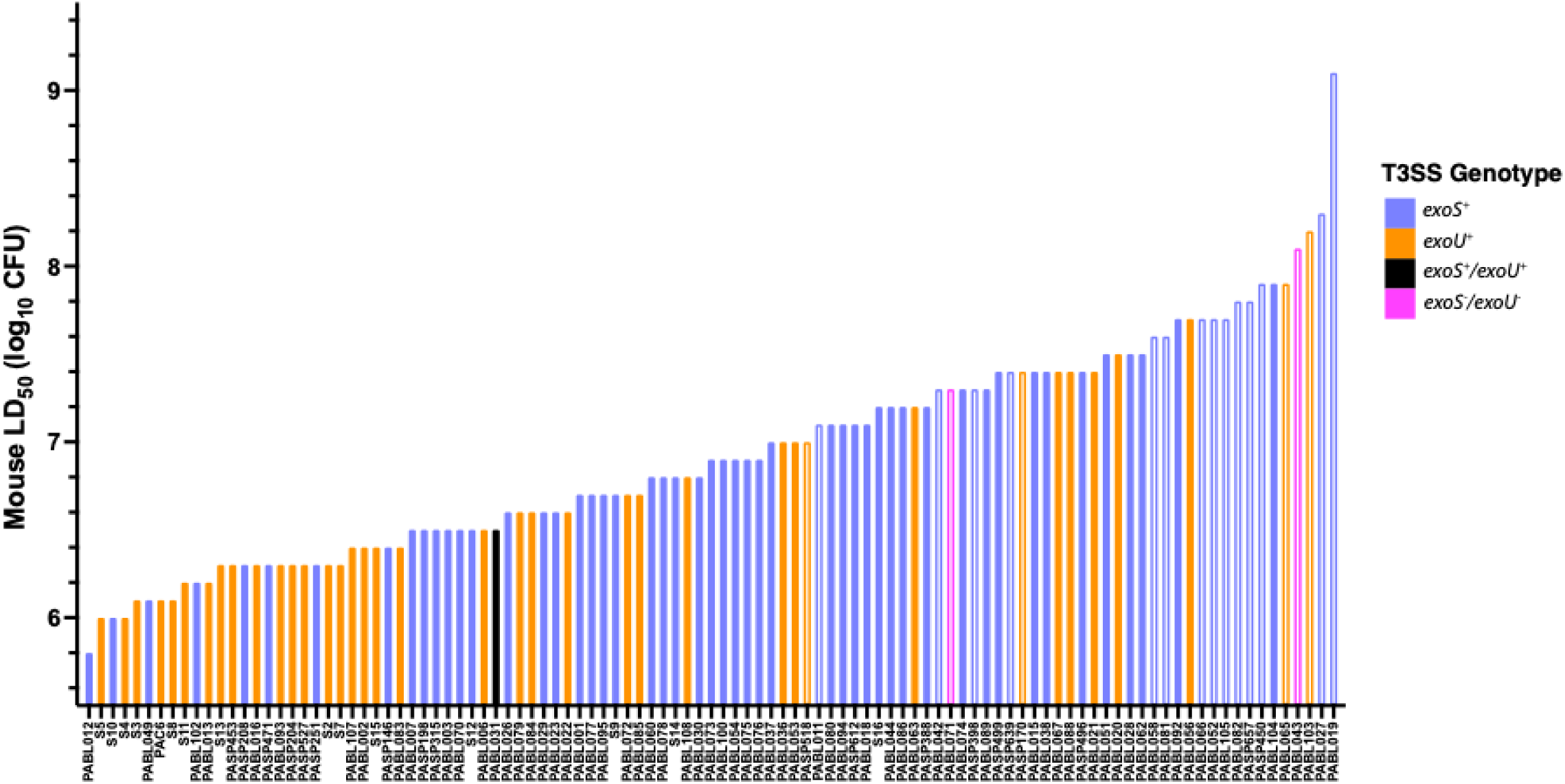
Virulence of *P. aeruginosa* clinical isolates in mice. BALB/c mice were previously challenged intravenously by tail vein injection with varying doses of the 105 *P. aeruginosa* clinical isolates and monitored for the development of pre-lethal infections. Results were used to calculate LD_50_ values for each isolate. T3SS effector genotype for each isolate is indicated by bar color. Bar fill indicates detection of *in vitro* effector secretion (solid; either ExoS^+^ or ExoU^+^) or lack thereof (open, ExoS^-^/ExoU^-^). For convenience, the LD_50_ values for all isolates except PABL019 were obtained and reformatted from Table S3 of Pincus et al. [32], distributed under the terms of the Creative Commons Attribution 4.0 International license (https://creativecommons.org/licenses/by/4.0/). For PABL019, the mouse LD_50_ value was obtained from Allen et al. [40].

### Virulence in a *G. mellonella* infection model

We next quantified the virulence of each isolate in *G. mellonella*. Unlike other bacterial species, many *P. aeruginosa* strains kill *G. mellonella* at extremely low inocula, where LD_50_ estimates become difficult to obtain [10]. We therefore used time to death as a measure of virulence, as we have previously shown this approach provides a consistent and reliable measure of *G. mellonella* virulence [10]. Isolate S7 was the most virulent, with an LT_50_ value of 10.3 hours (Fig. 3). PABL019 was the least virulent isolate, with an LT_50_ value of 35.6 hours. The median LT_50_ value for all isolates was 12.6 hours. These results demonstrate substantial variability in the virulence of *P. aeruginosa* isolates in the *G. mellonella* model.

**Figure 3.**
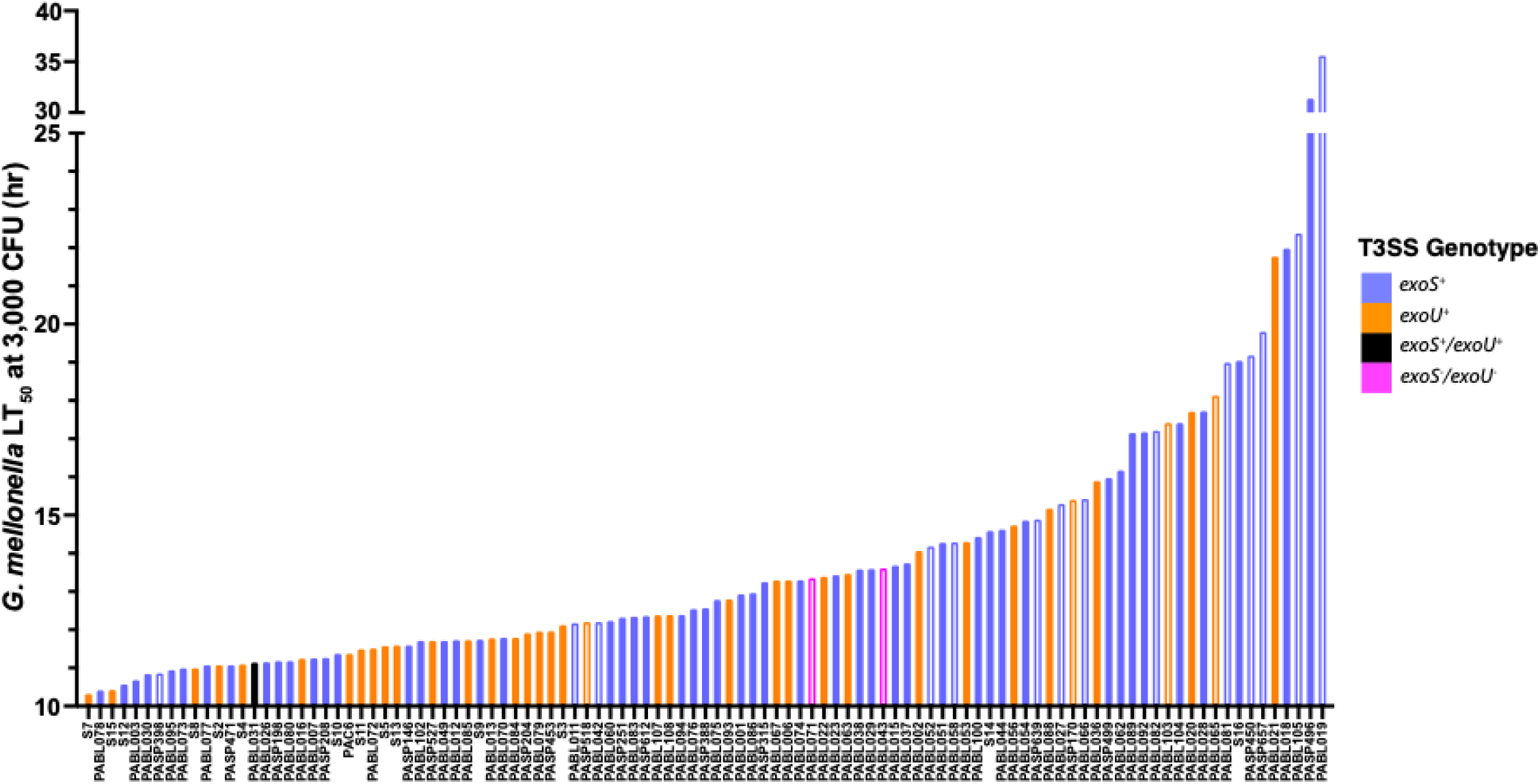
Virulence of *P. aeruginosa* clinical isolates in *G. mellonella*. *G. mellonella* larvae were infected with each *P. aeruginosa* isolate, and virulence was quantified by extrapolating an LT_50_ value for a dose of 3,000 CFU. T3SS effector genotype for each isolate is indicated by bar color. Bar fill indicates detection of *in vitro* effector secretion (solid; either ExoS^+^ or ExoU^+^) or lack thereof (open, ExoS^-^/ExoU^-^).

### Virulence in a cell co-culture system

Next, each isolate was tested for its ability to lyse human cells. We chose A549 cells, which are a cell line of pulmonary epithelial-like cells, because the lungs are a common site of *P. aeruginosa* infections [48] and epithelial cells have been used extensively to characterize *P. aeruginosa* cytotoxicity [15,49–51]. Bacteria were co-incubated with A549 cells, and cytotoxicity was quantified after 3 hours, a time point commonly used to measure *P. aeruginosa* cytotoxicity [15,52]. Most of the isolates caused little or no cytotoxicity, but 32 (30%) lysed greater than 5% of the A549 cells (Fig. 4, Supplemental Data). Among these 32 isolates, PABL072 showed the highest cytotoxicity (52%), and the average cytotoxicity for this subset of isolates was 33%. Thus, only a minority of isolates caused appreciable cytotoxicity under these conditions.

**Figure 4.**
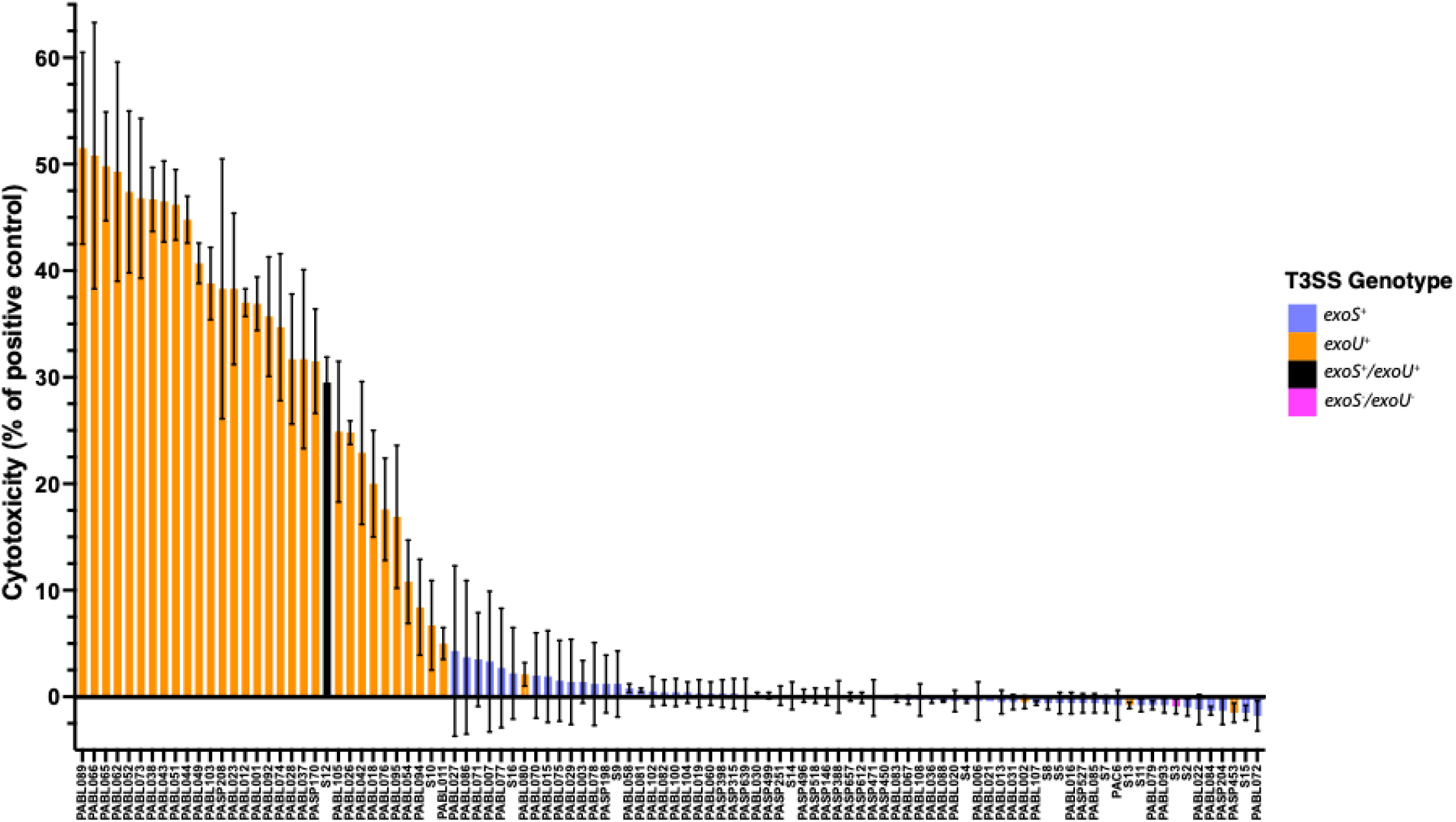
Virulence of *P. aeruginosa* clinical isolates in an epithelial-like cell lysis model. A549 cells were co-incubated with each isolate for 3 hr, and lactate dehydrogenase (LDH) was measured as a proxy for cell lysis. Detergent-induced lysis served as the positive control for complete lysis. T3SS effector genotype for each isolate is indicated by bar color. Bar fill indicates detection of *in vitro* effector secretion (solid; either ExoS^+^ or ExoU^+^) or lack thereof (open, ExoS^-^/ExoU^-^). Bars indicate the mean of three biological replicates; error bars indicate standard deviations.

### Comparison of *G. mellonella* and cell co-culture models to the mouse model

We compared the virulence measures of the 105 *P. aeruginosa* isolates in the *G. mellonella* and cell co-culture models to those of the mouse bacteremia model. To avoid the assumption of a linear relationship between virulence measurements across models, we used Spearman’s rank correlation coefficient (ρ) to quantify monotonic associations. Extrapolated LT_50_ values for 3,000 CFU in *G. mellonella* correlated in the “semistrong” range when compared to mouse LD_50_ values (ρ = 0.75, *P* < 0.0001, Fig. 5A). Similar correlations were observed using LT_50_ values for 300 CFU (ρ = 0.75, *P* < 0.0001) and 30,000 CFU (ρ = 0.64, *P* < 0.0001) (Supplemental Fig. 2). In contrast, comparison of cell cytotoxicity to mouse LD_50_ values yielded a correlation coefficient in the “weak” range (ρ = -0.47, *P* < 0.0001, Fig. 5B). This was largely driven by the absence of appreciable cytotoxicity associated with two-thirds of the isolates, preventing this model from discriminating virulence differences among these isolates. Thus *G. mellonella* was superior to the cell co-incubation model in measuring *P. aeruginosa* virulence when a mouse model of bacteremia was used as the gold standard.

**Figure 5.**
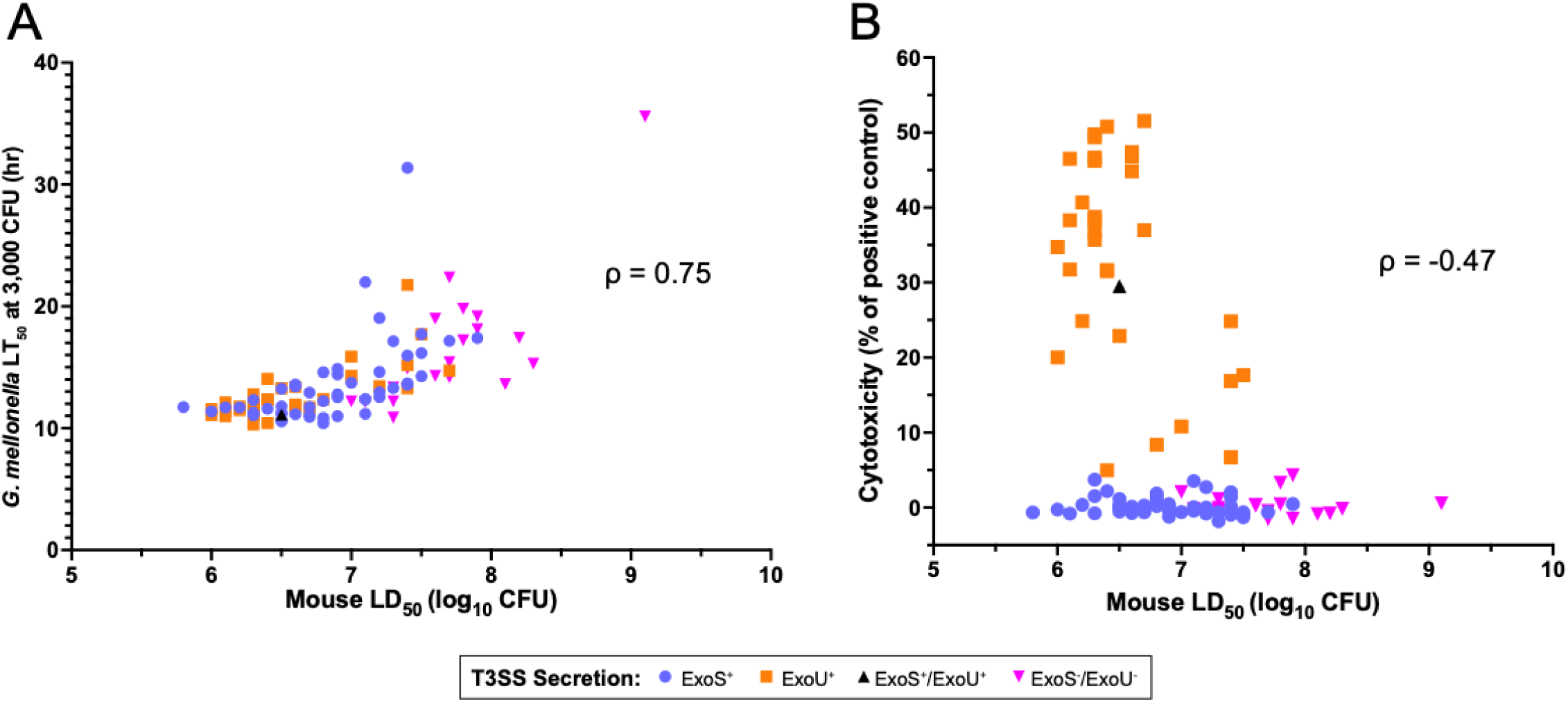
Concordance of *G. mellonella* or cell co-culture cytotoxicity with mouse virulence measurements. (A) The LT_50_ values at an extrapolated dose of 3,000 CFU in *G. mellonella* are plotted against corresponding LD_50_ values from the murine bacteremia model, yielding a Spearman’s rank correlation coefficient (ρ) of 0.75 (*P* < 0.0001). (B) A549 cytotoxicity values are plotted against corresponding LD_50_ values from the murine bacteremia model, yielding a ρ of -0.47 (*P* < 0.0001).

One potential difficulty in comparing the three model systems is that their readouts differ in range. While mouse LD_50_ values vary markedly, necessitating the use of a log scale, virulence in *G. mellonella* and cell culture systems vary less and are suitable for a linear scale. To better facilitate comparisons across these ranges, isolates were ranked from highest (rank 1) to lowest (rank 105) virulence for each model. A heat map was then generated to visually compare the relative virulence of isolates across models (Fig. 6). Isolates with a high mouse virulence rank (1-35) were significantly more likely to have a high *G. mellonella* virulence rank (Fisher’s exact test, *P* = 1.12 x 10⁻⁷), and isolates with a low virulence rank in mice (70-105) were also more likely to have a low virulence rank *G. mellonella* (*P* = 5.42 x 10^-9^). In contrast, isolates within the intermediate mouse virulence rank (36-69) were not significantly more likely to have an intermediate *G. mellonella* virulence rank (*P* = 0.12). For example, moderately virulent isolates in mice such as PABL078, PABL030, and PABL073 (ranks 49, 52, and 53, respectively) were among the most virulent in *G. mellonella* (ranks 2, 6, and 9, respectively). In contrast, cell co-culture rankings correlated less well with those of the mouse model. Although isolates with a high mouse virulence rank were significantly more likely to have a low cell survival rank (i.e., high cytotoxicity; *P* = 2.04 x 10-5), there were some highly virulent isolates (e.g., PABL012, S10, PABL049) that exhibited very little cytotoxicity. Isolates with low mouse virulence ranks did not show a significant association with high cell survival ranks (*P* = 0.051). These observations suggest that *G. mellonella* adequately identifies isolates that are at the extremes of virulence in mice but that cell co-culture cytotoxicity measurements do so less well.

**Figure 6.**
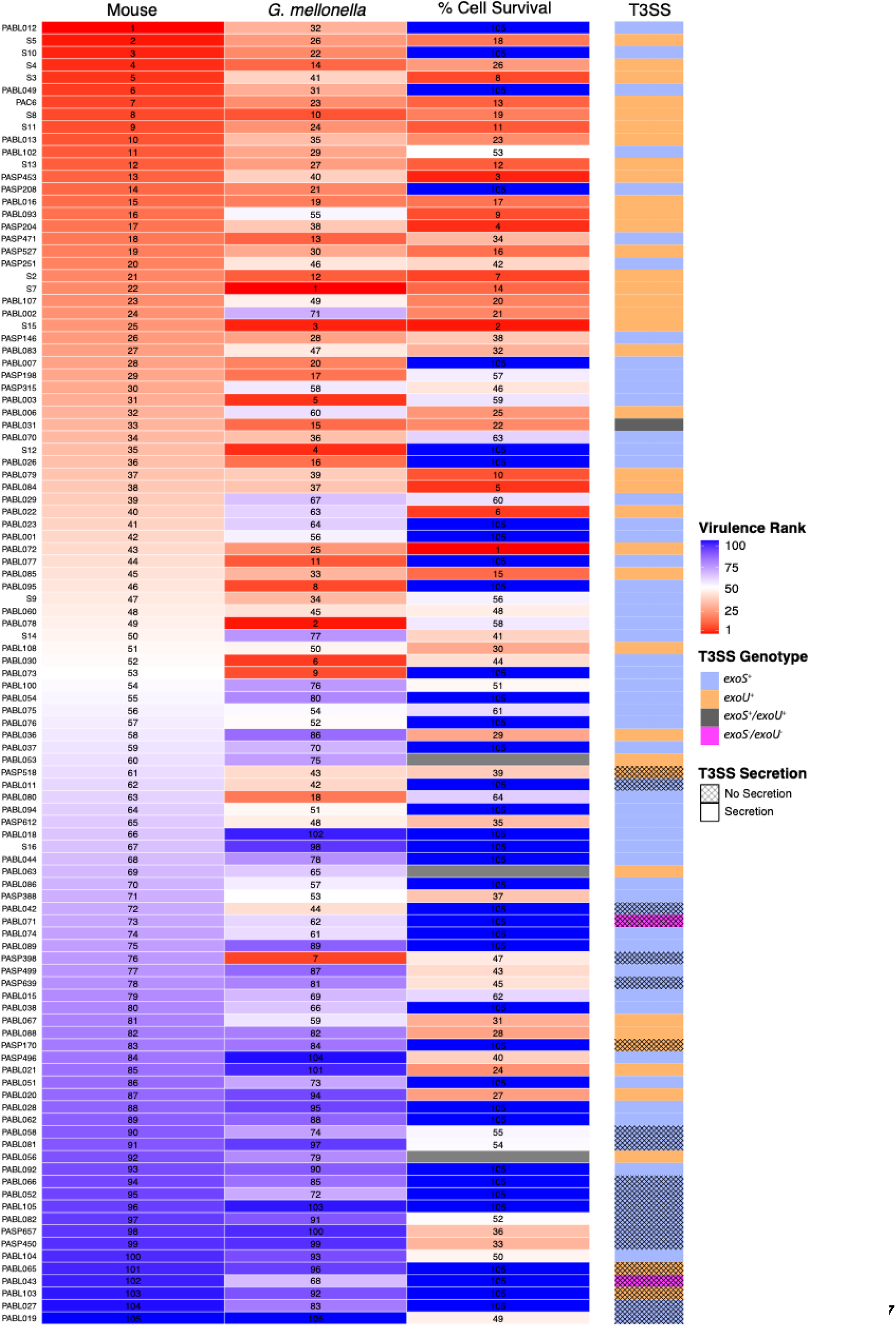
Ranked comparison of the virulence of the *P. aeruginosa* isolates measured by the three infection models. Mouse LD_50_ values (“Mouse”), *G. mellonella* LT_50_ values at 3,000 CFU (“*G. mellonella*”), and A549 percent cell survival (derived from cytotoxicity measurements; “% Cell Survival”) are shown in a heatmap format. For each metric, isolates are ranked from 1 (most virulent by that metric) to 105 (least virulent), and tiles are colored by virulence rank (see scale). In general, when multiple isolates shared identical virulence values within the same infection model isolates were assigned consecutive ranks in an arbitrary order. However, for “% Cell Survival,” data, where many isolates exhibited 100% cell survival, all isolates with 100% survival were ranked “105.” Isolates are ordered based upon their virulence rank in mice. The adjacent annotation bar (“T3SS”) indicates each isolate’s T3SS effector genotype (by color) and secretion phenotype (by crosshatch pattern). In the text, an isolate that secretes neither ExoS nor ExoU is referred to as ExoS^-^/ExoU^-^.

### The usefulness of *G. mellonella* to address population-based questions of *P. aeruginosa* virulence

Since *G. mellonella* infection outcomes had a semistrong correlation with those in mice, we next asked whether this insect model could be used to answer questions about *P. aeruginosa* pathogenesis. We used the “gold standard” mouse virulence values to answer the question and then determined whether the *G. mellonella* model provided the same answer.

We first asked whether antibiotic resistant isolates were less virulent than antibiotic susceptible isolates. The minimum inhibitory concentrations (MICs) of six antibiotics, each representing a major class of antipseudomonal antibiotics, were determined for all isolates (Supplemental Data). Isolates were then ranked from 0 to 6 based on the number of tested antibiotics to which they exhibited non-susceptibility. The most commonly observed non-susceptibility was to aztreonam (48%) and the least was meropenem (18%). As a group, isolates with higher ranks of antibiotic resistance exhibited significantly higher mouse LD_50_ values (lower virulence) (Kendall’s rank correlation, τ = 0.20, *P =* 0.007) (Fig. 7A), confirming previous reports [27,34–37]. This relationship between antibiotic non-susceptibility and virulence was also detected by the *G. mellonella* model (τ = 0.30, *P =* 4.9×10^-5^) (Fig. 7B). These findings suggest that the *G. mellonella* model is adequate to uncover relationships between virulence and antibiotic resistance.

**Figure 7.**
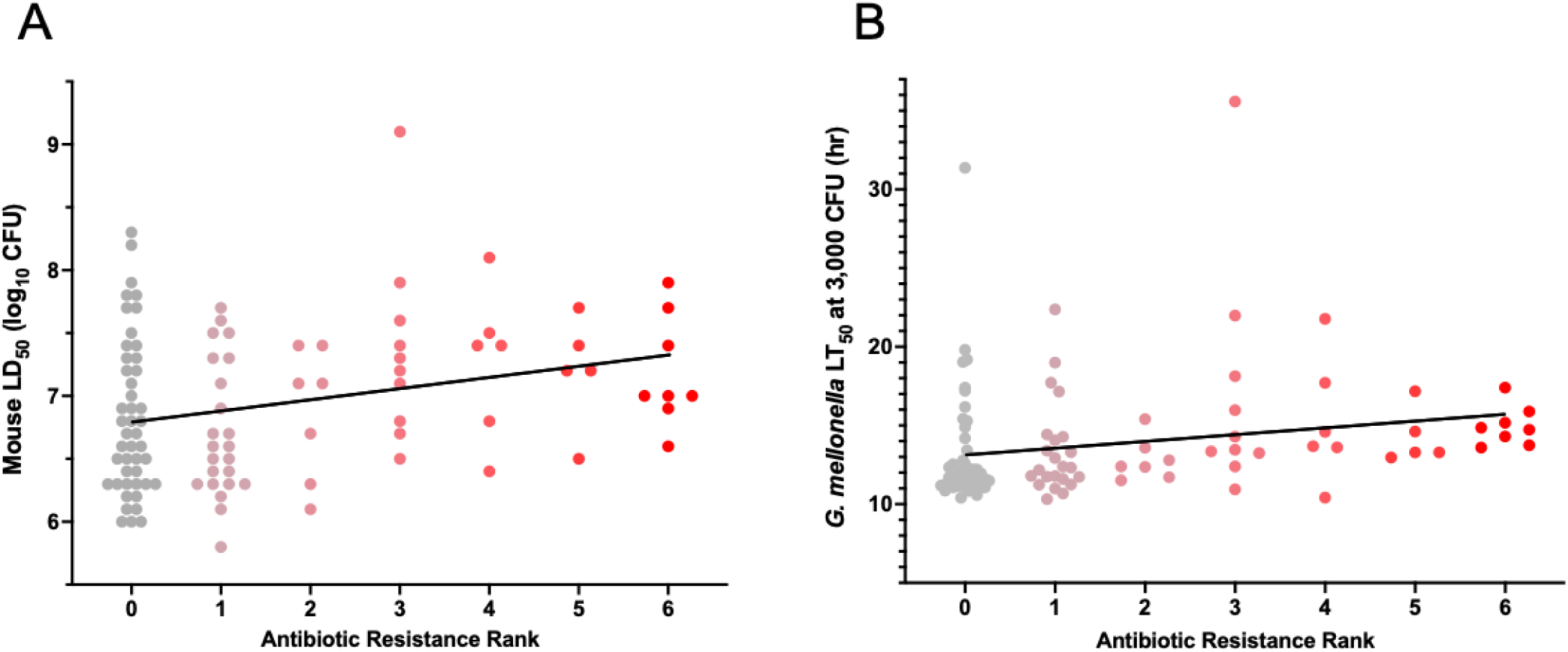
Relationship between antibiotic resistance and virulence of *P. aeruginosa* isolates in mice and *G. mellonella*. Each isolate was assigned an antibiotic resistance rank (0-6) based on the number of six tested antibiotics to which it was non-susceptible (i.e., intermediately susceptible or resistant). (A) Comparison of antibiotic resistance ranks to mouse LD_50_ values. A modest positive association between antibiotic-resistance levels and mouse LD_50_ values was observed (Kendall’s rank correlation, τ = 0.20, *P =* 0.007). Kendall’s tau (τ) indicates the strength and direction of association between two variables (e.g., τ = 1 indicates a perfect positive association, and τ = 0 indicates no association). (B) Comparison of antibiotic resistance ranks to *G. mellonella* LT_50_ values. A similar degree of association (τ = 0.30, *P =* 4.9×10^-5^) was noted. Each symbol represents an individual *P. aeruginosa* isolate.

We next asked whether some lineages of *P. aeruginosa* were characterized by unusually high levels of virulence, similar to what has been observed with *Staphylococcus aureus* [53], *Listeria monocytogenes* [54], and *Klebsiella pneumoniae* [55]. We first focused on HRCs due to their role in causing disproportionate numbers of infections. Currently, consensus criteria for defining lineages as HRCs have not yet emerged, and different definitions have been used [25,26,29]. For this study, we defined *P. aeruginosa* HRCs as STs reported in the literature as “HRCs”, found on at least two continents, reported to be highly resistant to antibiotics, and more frequently cultured from patients than other *P. aeruginosa* lineages. To address the last criterion, we determined the relative abundance of each ST among *P. aeruginosa* genomes deposited in NCBI (Supplemental Fig. 3). Using these criteria, we identified 10 STs that represented the most common HRCs present in our collection. A total of 32 isolates (30.5%) in our collection belonged to these 10 HRC STs. The virulence of these HRCs did not differ significantly from non-HRCs in either the mouse or *G. mellonella* infection model (Fig. 8A and 8B). These findings remained consistent when the analysis was restricted to the five most common HRCs in the NCBI repository (Fig. 8C and 8D). Interestingly, our analysis demonstrated that many of the lineages defined as HRCs in the literature contain a relatively low proportion of antibiotic-resistant isolates (Fig. 8). Together, these findings suggest that HRCs are no more or less virulent than non-HRC *P. aeruginosa* lineages, and that the *G. mellonella* model mimics the mouse model in demonstrating this relationship.

**Figure 8.**
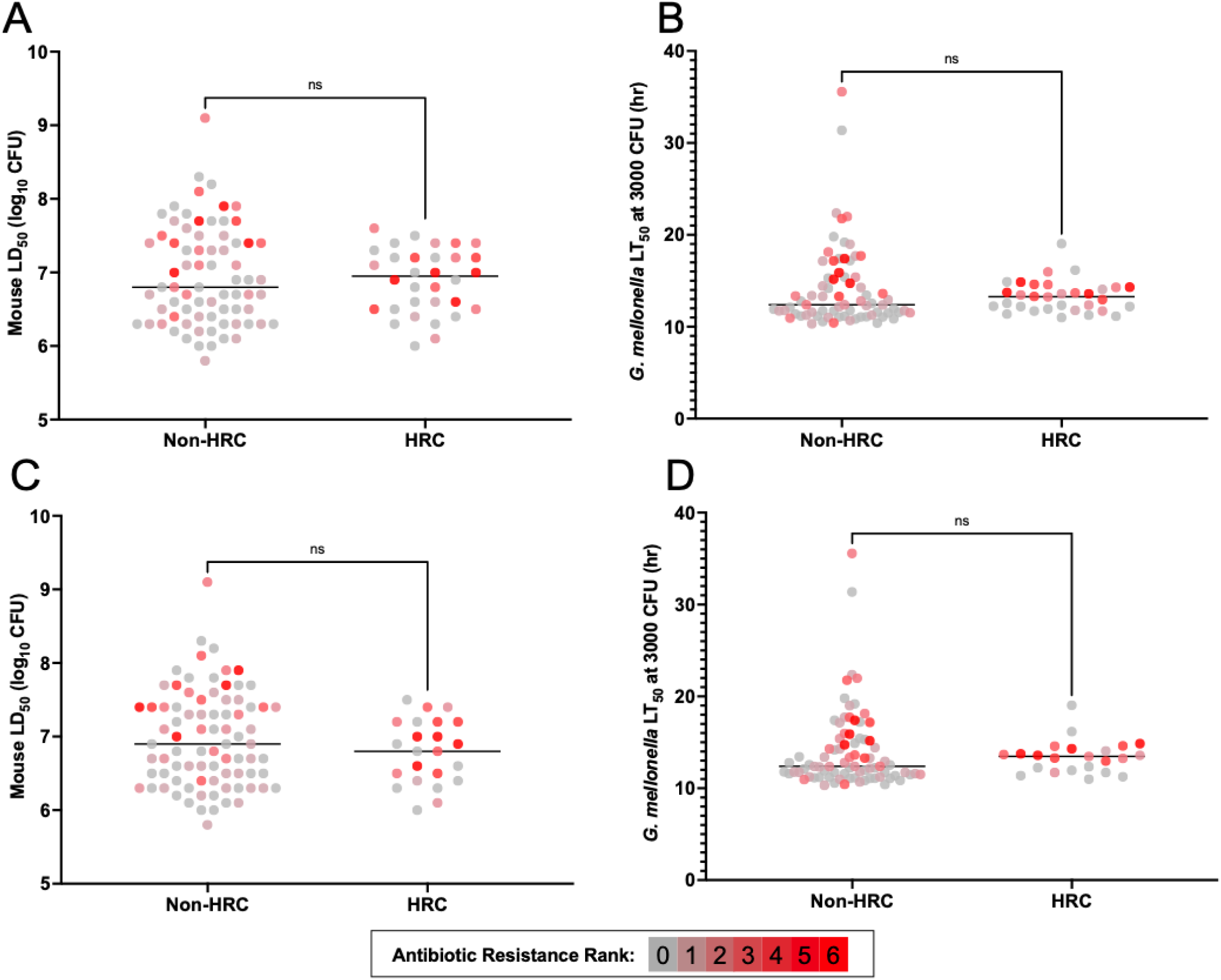
Relationship between HRC status and virulence in mice and *G. mellonella*. Comparison of (A) mouse LD_50_ values and (B) *G. mellonella* LT_50_ values of isolates belonging to 10 HRC STs (235, 111, 274, 253, 244, 395, 179, 308, 175, 463) to those of the remaining isolates. No statistical differences between the two groups were noted (*P* = 0.90 and *P* = 0.46, respectively). Comparison of (C) mouse LD_50_ values and (D) *G. mellonella* LT_50_ values of isolates belonging to five HRC STs (235, 111, 274, 253, 244) to those of the remaining isolates. Again, no statistical differences were noted (*P* = 0.31 and *P* = 0.43, respectively). Each symbol represents an individual isolate. The antibiotic resistance rank (0 to 6) of each isolate is indicated by gray to red hue. Mann-Whitney U test, ns = not significant.

We next examined whether any lineages of *P. aeruginosa* are associated with unusually high levels of virulence, independent of HRC status. We grouped isolates based on core genome similarity using a phylogeny-based clustering tool (Supplemental Fig. 4) [56]. Seven clusters of genetically related isolates were generated, with five isolates not clustering into any group (Fig. 9). Mouse LD_50_ values differed significantly across clusters (Kruskal-Wallis test, *P* = 0.006). Cluster 7, comprised of all *exoU*^+^ isolates and PABL071 (*exoS^-^*/*exoU*^-^), had the lowest median LD_50_ value (highest virulence), but no cluster was significantly more or less virulent by pairwise comparisons. Similarly, the clusters did not significantly differ in their virulence in *G. mellonella.* This analysis did not identify hypervirulent lineages of *P. aeruginosa*, suggesting that either such lineages do not exist or that analysis of larger isolate collections allowing for more granular groupings will be necessary to identify such lineages. In addition, the mouse and *G. mellonella* models yielded similar results when applied to this question.

**Figure 9.**
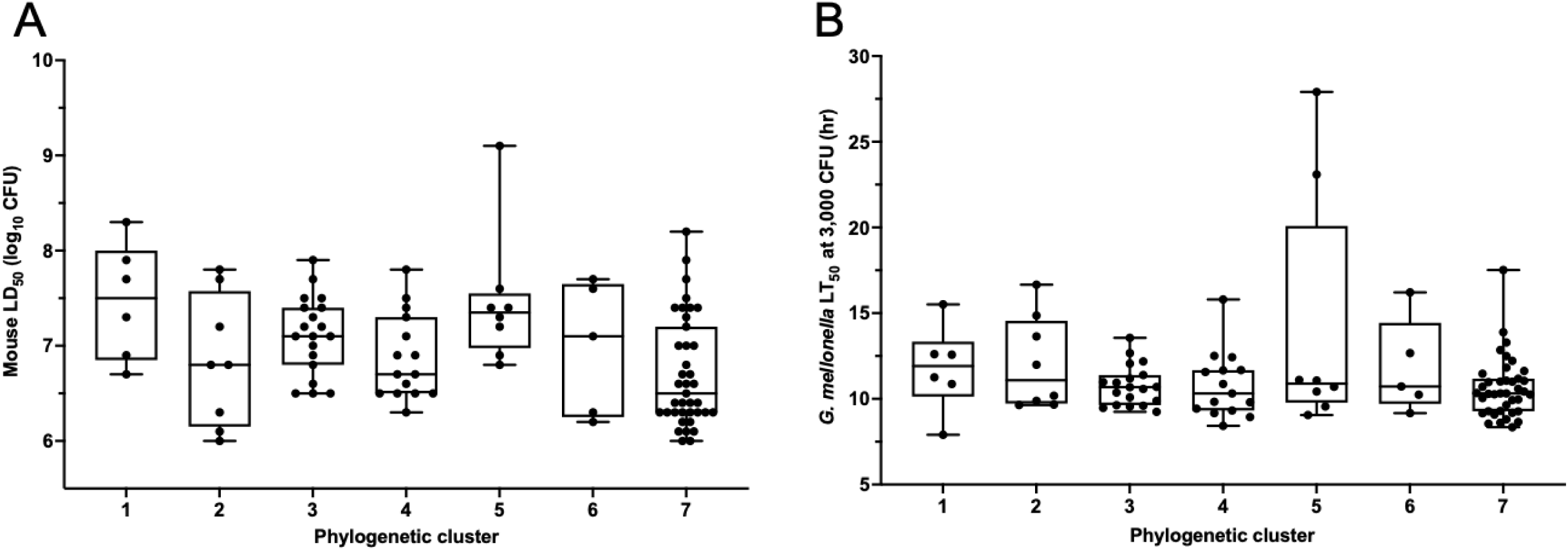
Comparison of the virulence of *P. aeruginosa* isolates in distinct genetic clusters. (A) Comparison of the mouse LD_50_ values of isolates belonging to seven defined phylogenetic clusters. Median mouse LD_50_ values differed significantly across clusters (Kruskal–Wallis test*, H* = 17.92, *P* = 0.006), but no significant pairwise differences were noted (Bonferroni-adjusted Dunn’s post-hoc multiple comparisons, *P* > 0.05). (B) Comparison of *G. mellonella* LT_50_ values of isolates belonging to the seven clusters. No significant differences were found between the clusters (*H* = 4.69, *P* = 0.58). Each symbol represents an individual isolate. Bars represent medians, boxes represent the interquartile range (25th–75th percentile), and brackets represent minimum and maximum values.

### Differential roles of the T3SS in the models

To explore the mechanistic basis for the differential performance of the mouse and *G. mellonella* models, we focused on the T3SS, as it is a major pathogenic determinant of *P. aeruginosa* and has been shown to impact virulence in both models [23,57].

In mice, ExoU^+^ and ExoS^+^ isolates were generally more virulent than ExoS^-^ /ExoU^-^ isolates (*P* < 0.0001), with 11 of the 12 least virulent isolates being ExoS**^-^/**ExoU**^-^**(Fig. 2). As a group, ExoU^+^ isolates were more virulent than ExoS**^+^** isolates (*P* = 0.041, Fig. 10A), with 18 of the 25 most virulent isolates being ExoU^+^ (Fig. 2). The overall population also displayed a moderate phylogenetic signal associated with virulence (Pagel’s λ = 0.60, *P* = 4.79 x 10^-5^, Supplemental Fig. 5A), which may be partly driven by the phylogenetic clustering of *exoU^+^* Group B isolates. However, several ExoS^+^ isolates exceeded ExoU^+^ isolates in virulence, suggesting that additional factors beyond these T3SS effector proteins contribute to overall virulence. In fact, the most virulent isolate overall was PABL012, an ExoS^+^ isolate (Fig. 2). These results are consistent with previously observed relationships between T3SS phenotypes and disease severity in mice and in patients [15,20,21,23].

**Figure 10.**
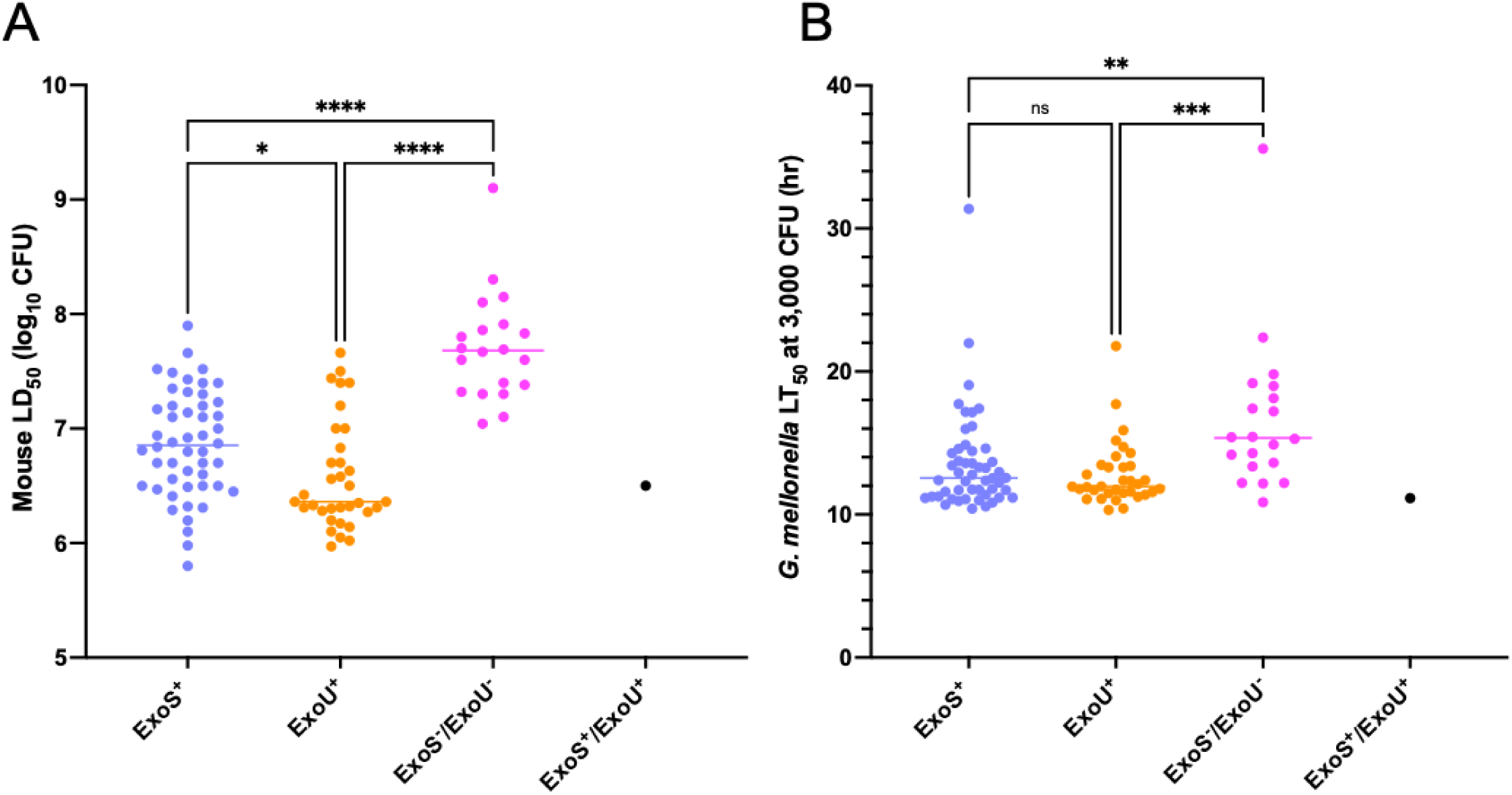
Comparison of virulence of *P. aeruginosa* isolates in mice and *G. mellonella* based on ExoS and ExoU secretion status. (A) Mouse virulence data for each T3SS phenotype. ExoS^+^ isolates were less virulent than ExoU^+^ isolates (*P* = 0.041). (B) *G. mellonella* virulence data for each T3SS phenotype. No significant difference in virulence between ExoS^+^ and ExoU^+^ isolates was observed (*P* = >0.99). Each symbol represents an individual *P. aeruginosa* isolate. Kruskal-Wallis test; \**P* <0.05, \*\**P* < 0.01, \*\*\**P* < 0.001, \*\*\*\**P* < 0.0001, ns = not significant.

Results using *G. mellonella* were similar to those in mice but with a few notable differences. Both ExoS^+^ and ExoU^+^ isolates were more virulent than ExoS**^-^/**ExoU**^-^**isolates (*P* = 0.0034 and *P* = 0.0006, respectively), but there was substantial overlap between all three groups (Fig. 10B). For example, the seventh most virulent isolate in *G. mellonella* was PASP398, an ExoS**^-^/**ExoU**^-^** isolate (Fig. 3). In contrast to the mouse model, ExoU^+^ isolates, as a group, were not significantly more virulent than ExoS^+^ isolates (Fig. 10B); 6 of the 10 most virulent isolates were ExoS^+^ (Fig. 3). Additionally, the population displayed no evidence of any phylogenetic signal associated with virulence (Pagel’s λ = 0.008, *P* = 0.87, Supplemental Fig. 5B), suggesting virulence is not lineage specific in this model. One explanation for these results is that individual T3SS effector proteins have differential impacts on virulence in *G. mellonella* and mice, which may partially explain why the correlation between virulence measures in these two models was not higher.

Several isolates exhibited pronounced discordance in virulence between the mouse and *G. mellonella* models (Fig. 6). We assessed whether ExoU and ExoS secretion phenotypes contribute to these model-dependent differences by focusing on isolates with the greatest divergence in virulence ranks between models. We identified the 10 isolates for which the virulence rank in *G. mellonella* most exceeded the virulence rank in mice (i.e., were relatively more virulent in *G. mellonella* than in mice) and the 10 isolates for which the virulence rank in mice most exceeded that in *G. mellonella* (i.e., were relatively more virulent in mice than in *G. mellonella*) (Fig. 6, Supplemental Table 1). Both sets of isolates spanned a large range of virulence values in each model (Supplemental Fig. 6). However, isolates with a greater relative virulence rank in *G. mellonella* were statistically more likely to be from Group A (predominantly *exoS^+^* isolates) than were those with a greater relative virulence rank in mice (*P* = 0.033, Table 1, Supplemental Fig. 7). This finding, together with the observation that ExoS^+^ isolates had higher virulence relative to ExoU^+^ isolates in *G. mellonella* than in mice (Fig. 10), suggests that *G. mellonella* are more affected by ExoS or another Group A-associated factor than mice.

**Table 1.** Phylogenetic clade distribution among isolates with the greatest relative virulence rank difference between the mouse and *G. mellonella* models.

|  | Number of isolates with the greatest relative virulence rank<sup>a</sup> |  |
| --- | --- | --- |
| Phylogenetic Clade | *G. mellonella* | Mouse |
| Group A | 9 | 5 |
| Group B | 0 | 5 |
| Fisher's exact test, *P* = 0.033 |  |  |

<sup>a</sup>Top 10 isolates for which the virulence rank in *G. mellonella* exceeded that in mice, and the top 10 isolates for which the virulence rank in mice exceeded that in *G. mellonella*. PABL043 (a PA7-like isolate) was excluded from this analysis because it belongs to neither Group A or Group B.

We next examined whether *in vitro* type III secretion might explain why the cell co-culture results differed from the mouse results. This difference was largely driven by the fact that the ExoS^+^ and ExoS^-^/ExoU^-^ isolates all displayed <5% cytotoxicity despite showing a broad range of mouse LD_50_ values (Fig. 4, Supplemental Data). In contrast, all ExoU^+^ isolates caused appreciable cytotoxicity, consistent with previous reports demonstrating ExoU is the predominant cytotoxin of *P. aeruginosa* [14,58]. Substantial differences were apparent in the proportion of cells lysed by ExoU^+^ isolates during infection, suggesting that factors other than or in addition to ExoU are involved in modulating the cytotoxicity of these isolates, potentially including differences in *exoU* expression or effector delivery. However, even when only ExoU^+^ isolates were included in the analysis, cytotoxicity correlated poorly with mouse LD_50_ values (ρ = -0.30, *P* = 0.095), indicating that cytotoxicity alone only modestly predicts virulence in the murine model (Supplemental Fig. 8).

## Discussion

To test the utility of using *G. mellonella* and cell co-culture models as alternatives to a mouse model of bacteremia, we quantified the virulence of *P. aeruginosa.* We found that *G. mellonella* produced results comparable to those of the mouse model but cell lysis of A549 epithelial-like cells was a less reliable measure. We then showed that *G. mellonella* and mice yielded similar answers to *P. aeruginosa* population-based questions regarding virulence. These findings suggest that *G. mellonella* provides a high-throughput, cost-effective and scalable substitute for the mouse bacteremia model in screening large numbers of *P. aeruginosa* isolates for virulence. However, final conclusions should be confirmed in mice.

Our results support previous reports showing a correlation between *P. aeruginosa* virulence measured by *G. mellonella* and murine models [10–12] and further quantify the extent of this correlation. Several biological differences may explain why the correlation was not higher. Mice and *G. mellonella* are markedly different in anatomy, physiology, and immunology. In addition, the method of inoculation (parenchymal injection in *G. mellonella* vs. intravenous injection in mice) differed, as likely did dissemination [59,60]. Additionally, *P. aeruginosa* virulence factors may be differentially active in each host, either from variations in their targets or in host defense systems that detect or neutralize their activities. In this regard, a closer examination of our results provides several clues to these molecular differences. First, ExoS^+^ isolates as a group were as virulent as ExoU^+^ isolates in *G. mellonella* but less virulent in mice. For mice, this observation is consistent with previous reports demonstrating that in a mouse pneumonia model deletion of *exoU* attenuated virulence to a greater degree than deletion of *exoS* [23]. It is possible that ExoS or another Group A-associated factor is more active in *G. mellonella* than in mice or that ExoU is less active. Notably, the role of ExoS in *G. mellonella* killing has not been evaluated, but an *exoU*-deficient mutant of *P. aeruginosa* was not attenuated in killing *G. mellonella* [57]. Second, ExoS^-^/ExoU^-^isolates were generally less virulent in mice, but some of these isolates exhibited high virulence in *G. mellonella*, such as PASP398, PABL042, and PABL011 (Fig. 6). This finding suggests that unidentified factors other than ExoS and ExoU contribute relatively more to outcomes in *G. mellonella* than in mice. Defining which *P. aeruginosa* virulence determinants differentially impact illness severity in mice and *G. mellonella* is an exciting new avenue of research that may better define the conditions under which *G. mellonella* can be used with confidence as a substitute for mice.

Several isolates in our collection warrant further investigation. The first is PABL019, which was the least virulent isolate in both *G. mellonella* and mice (rank #105, Figs. 5A and 6). Interestingly, this isolate was obtained from a blood culture of an individual with CF. CF-associated *P. aeruginosa* isolates have been reported to be attenuated in virulence in both models [12,61], likely due to their adaptation during chronic infection of the airways [3,62–64]. Nearly all of the 105 tested isolates in our study were from acute infections, so we were unable to further examine the behavior of CF isolates in *G. mellonella,* but future studies using such isolates would further test the ability of *G. mellonella* to mirror mice in this context. The second isolate is PABL002, which demonstrated high virulence in mice (rank #24) but relatively modest virulence in *G. mellonella* (rank #71) (Supplemental Table 1). This isolate may be useful in defining those pathogenic factors that are critical in mice but fail to influence virulence in *G. mellonella*. Conversely, PASP398 had low virulence in mice (rank #76) and relatively high virulence in *G. mellonella* (rank #7), suggesting the presence of factors that enhance pathogenicity in *G. mellonella* but not in mice. These and similar isolates may serve as useful tools to dissect model-specific virulence determinants.

In comparison to *G. mellonella,* the cell co-culture model performed less well in predicting virulence in mice (Fig. 5B). At first glance, this result is surprising, as ExoU^+^ isolates exhibit high levels of virulence in mice and are also highly cytotoxic against epithelial cell lines (Fig. 2) [13–15,23]. However, closer examination suggests three contributors to the weaker correlation between these models. First, although ExoU^+^ isolates exhibited notable cytotoxicity, the remaining isolates caused minimal cytotoxicity. As a result, this assay was unable to discriminate virulence differences within the groups of ExoS^+^ and ExoS^-^/ExoU^-^ isolates, which comprised ∼67% of the tested isolates. Second, while many ExoU^+^ isolates were indeed highly virulent in mice and associated with substantial cell lysis, many noncytotoxic ExoS^+^ isolates were also highly virulent in the mouse model, such as PABL012, S10, and PABL049 (Fig. 6). Third, even among ExoU^+^ isolates, the correlation between cytotoxicity and mouse LD_50_ values was poor (ρ = -0.30, Supplemental Figure 8). *P. aeruginosa* harbors multifactorial mechanisms to disseminate in the host, evade the immune response, and alter host physiology; epithelial cells alone may not capture the complexity of these host-pathogen interactions [65]. Despite the well-established cytotoxic effects of ExoU, its contribution to overall virulence may be modulated or supplemented by additional factors. Alternatively, cell lysis may be an *in vitro* phenomenon that does not reflect a more nuanced pathogenic role for ExoU *in vivo.* Finally, the conditions of our cell lysis assays, such as the MOI, time of co-incubation, and host cell type may not reflect the *in vivo* conditions under which ExoU exerts its impact on disease progression [66].

The impact of antimicrobial resistance on *P. aeruginosa* virulence is somewhat controversial. Our findings with 105 *P. aeruginosa* isolates across two infection models show an inverse relationship between antibiotic resistance and virulence, which could indicate a tradeoff between these two properties. However, these associations do not necessarily imply that antimicrobial resistance determinants themselves cause reduced virulence. *P. aeruginosa* strains inhabiting hospitals and long-term care facilities are heavily exposed to antibiotics but also to other selective pressures intrinsic to survival in patients and the healthcare environment. Adaptations that facilitate persistence in these niches would be expected to co-occur with antibiotic resistance alleles and may ultimately be responsible for the inverse relationship we observed. For example, *P. aeruginosa* lineages cultured from chronically infected CF patients simultaneously accumulate disruptions of bacterial virulence attributes and mutations that enhance antibiotic resistance, presumably due to independent immune and antimicrobial selective pressures [64,67].

Hypervirulent clades have been well described in several bacterial species [53–55], but whether such clades exist in *P. aeruginosa* remains unclear. We were unable to detect groups of genetically related isolates that were highly virulent in mice or *G. mellonella*. In particular, the virulence levels of defined HRCs, which have previously been linked to poor clinical outcomes in patients [25–27,68], were not significantly different from non-HRCs in either model. This finding suggests that HRC-associated poor outcomes stem mainly from their resistance to antibiotic treatments or their propensity to infect patients with comorbid conditions. Interestingly, many isolates belonging to HRC STs in our collection were relatively antibiotic susceptible (Fig. 8). This finding supports previous reports suggesting that HRCs are more prone to acquire and maintain resistance determinants but that many isolates within HRCs remain susceptible to antibiotics [26].

Our study has several limitations. The clinical isolates we used predominantly originated from bloodstream infections and may not accurately reflect the virulence or clonal distribution of *P. aeruginosa* isolates from other infections. Additionally, sampling spanned only three continents; broader geographic coverage is necessary to demonstrate whether our results are generalizable. Unlike murine models, which use inbred mice, *G. mellonella* larvae may vary in genetic background and husbandry, introducing possible variability in LT_50_ values. Cytotoxicity was assessed exclusively in A549 cells, a well-established cell line for cytotoxicity assays, but results may differ with other cell lines. We used a 3-hr readout for cytotoxicity, but longer times likely would have detected cytotoxicity among non-ExoU isolates. However, this likely would come at the cost of saturated cell lysis for the ExoU^+^ isolates. Finally, we recognize that definitive conclusions on the utility of alternative infection models such as *G. mellonella* and cell co-culture assays cannot be drawn from a single type of mouse model (bacteremia) and cytotoxicity readout (cell lysis) with a single bacterial species (*P. aeruginosa*). However, we believe that this study will serve as an incremental step towards the accumulation of sufficient data to answer this important question

## Author Contributions

Conceptualization: A.V., A.R.H.

Formal analysis: A.V., C.A., T.J.K., J.H.

Funding acquisition: A.R.H., K.E.R.B.

Investigation: A.V., C.A., T.J.K., S.N., T.W., I.N., E.V., D.H., J.N., P.G., T.A., S.D.M., T.L.T., W.C., J.J.L., P.P., N.B.P., J.P.A., E.A.O., K.E.R.B.

Methodology: C.A., T.J.K., T.A.

Project administration: A.V., C.A., I.N., T.W.

Resources: A.O., C.-H.C., A.R.H.

Software: T.J.K.

Visualization: A.V., D.A.

Writing – original draft: A.V., A.R.H.

Writing – review & editing: A.V., A.R.H.

## Data Availability

All relevant data supporting the findings of this study are within the manuscript and its Supporting Information files. Whole-genome sequencing data for the isolates analyzed in this study were previously published (32, 40).

## Acknowledgements

We acknowledge the computational resources and staff support provided by the Genomics Compute Cluster, which is jointly supported by the Feinberg School of Medicine, the Center for Genetic Medicine, Feinberg’s Department of Biochemistry and Molecular Genetics, the Office of the Provost, the Office for Research, and Northwestern Information Technology. The Genomics Compute Cluster is part of Quest, Northwestern University’s high-performance computing facility, with the purpose of advancing research in genomics.

